# Acquisition of cross-azole tolerance and aneuploidy in *Candida albicans* strains evolved to posaconazole

**DOI:** 10.1101/2022.01.06.475277

**Authors:** Rebekah J. Kukurudz, Madison Chapel, Quinn Wonitowy, Abdul-Rahman Adamu Bukari, Brooke Sidney, Riley Sierhuis, Aleeza C. Gerstein

## Abstract

A number of *in vitro* studies have examined the acquisition of drug resistance to the triazole fluconazole, a first-line treatment for many Candida infections. Much less is known about posaconazole, a newer triazole. We conducted the first *in vitro* experimental evolution of replicates from eight diverse strains of *C. albicans* in a high level of the fungistatic drug posaconazole. Approximately half of the 132 evolved replicates survived 50 generations of evolution, biased towards some of the strain backgrounds. We found that although increases in drug resistance were rare, increases in drug tolerance (the slow growth of a subpopulation of cells in a level of drug above the resistance level) were common across strains. We also found that adaptation to posaconazole resulted in widespread cross-tolerance to other azole drugs. Widespread aneuploidy variation was also observed in evolved replicates from some strain backgrounds. Trisomy of chromosomes 3, 6, and R was identified in 11 of 12 whole-genome sequenced evolved SC5314 replicates. These findings document rampant evolved cross-tolerance among triazoles and highlight that increases in drug tolerance can evolve independently of drug resistance in a diversity of *C. albicans* strain backgrounds.

## INTRODUCTION

Drug resistance is a critical threat to global public health. Antimicrobial resistance is inherently an evolutionary phenomenon: drug-resistant individuals arise and spread within susceptible populations. The genetic basis and rate of adaptation in a drug is in part a deterministic process, akin to the evolutionary process under any environmental stress, influenced by the specifics of the environment, the microbial population size, the mutation rate, and the effect size of available beneficial mutations. Unlike bacteria, which frequently acquire plasmid-mediated beneficial genes and alleles from the environment, fungal microbes primarily adapt via vertical transmission. Genomic variation within evolving populations often includes small-scale point mutations and insertions and deletions as well as larger-scale karyotypic mutations in ploidy (the number of chromosome sets), aneuploidy (copy number change in one or several chromosomes), and zygosity (the number of alleles at a given position in the genome) (Selmecki *et al*. 2015; Ene *et al*. 2018, 2021; Wang *et al*. 2018). Few antifungal drug classes are currently approved for the treatment of fungal infections. One strategy to prolong the utility of existing drugs is to understand better the factors that influence fungal resistance acquisition, to reduce the likelihood that resistance will arise.

In addition to drug resistance, which is often measured as the minimum inhibitory concentration of drug that reduces growth by some amount (MIC, e.g., 50% or 80%) after 24 h, drug tolerance has recently emerged as an important parameter in characterizing drug response in fungal species. Fungal tolerance (which is distinct from bacterial tolerance (Levin-Reisman *et al*. 2017)) is defined as the proportion of the population that grows slowly in drug concentrations above the minimum inhibitory concentration (Rosenberg *et al*. 2018; Berman and Krysan 2020). Although few studies have quantified tolerance yet, it may play a role in persistent candidemia (Rosenberg *et al*. 2018) and mortality (Levinson *et al*. 2020). From an evolutionary perspective, beneficial mutations that arise in populations evolving in fungistatic environments that primarily inhibit rather than kill susceptible cells could act in two distinct pathways: they can increase resistance (i.e., increase the MIC), or they can increase tolerance (i.e., enable a larger proportion of the population to grow above the MIC). A recent screen of 235 clinical *Candida* spp. isolates found that resistant isolates also tended to be more tolerant to the fungistatic drug fluconazole (Salama & Gerstein, submitted). However, a large-scale *in vitro* evolution experiment in fluconazole found that changes in tolerance evolved independently of resistance (Gerstein and Berman 2020). As many antifungal drugs are fungistatic rather than fungicidal, increasing drug tolerance may be an unappreciated yet critical selective avenue for fungal populations adapting to a drug.

Relatively few species account for most human fungal infections, with *Candida albicans* the primary species responsible for mucosal disease, *Aspergillus fumigatus* for allergic disease, and *Trichophyton* spp. for skin infections (Bongomin *et al*. 2017). The azole drugs fluconazole and voriconazole are first-line treatments for candidiasis and aspergillosis, respectively (Pappas *et al*. 2016; Patterson *et al*. 2016). Primarily through the study of fluconazole-resistant *C. albicans* strains, the genetic basis of two major adaptive pathways have been identified: the first involves alterations or overexpression of *ERG11*, which produces the target demethylase (Asai *et al*. 1999; Lamb *et al*. 2000; Selmecki *et al*. 2008; Flowers *et al*. 2015; Paul *et al*. 2019; Lee *et al*. 2020); the second is through the up-regulation of drug efflux pumps encoded by *CDR1, CDR2, TAC1, MRR1*, and *MDR1* (Sanglard *et al*. 1995; Asai *et al*. 1999; Lamb *et al*. 2000; Coste *et al*. 2006; *Selmecki et al*. 2008; Flowers *et al*. 2015; Paul *et al*. 2019; Lee *et al*. 2020). One of the newest azoles, posaconazole, effectively treated infections resistant to first-line azoles in *A. fumigatus, C. albicans*, and *C. neoformans* (Xiao *et al*. 2004; Chau *et al*. 2004; Firinu *et al*. 2011; Sionov *et al*. 2012). Intriguingly, although specific point mutations in *ERG11* homologues confer posaconazole cross-resistance to azoles and other antifungal drugs in *Aspergillus* spp. (Lockhart *et al*. 2011; D’Agostino *et al*. 2018; Abastabar *et al*. 2019), single *C. albicans ERG11* point mutations do not provide the same degree of posaconazole cross-resistance (MacCallum *et al*. 2010; Sanglard and Coste 2016; Warrilow *et al*. 2019). Rather, it seems that multiple *ERG11* mutations are required for posaconazole resistance (Li *et al*. 2004). This may be attributable to the extended side chain of posaconazole interacting with an additional domain of the target enzyme (Li *et al*. 2004; Xiao *et al*. 2004; Chau *et al*. 2004; Katragkou *et al*. 2012).

*C. albicans* is a predominantly diploid asexual organism. Genetic diversity within populations is primarily achieved through mitosis, though parasexual reproduction is also possible (Hickman *et al*. 2013; Ene and Bennett 2014). In addition to point mutations, unicellular fungal microbes seem prone to acquiring chromosomal aneuploidies during adaptation (Gerstein and Berman 2015; Gerstein and Sharp 2021), and mitotic recombination results in loss-of-heterozygosity. Intriguingly, exposure to the triazole drugs fluconazole, ketoconazole, voriconazole, and itraconazole potentiate the appearance of chromosomal aneuploidy (Harrison *et al*. 2014). Extra copies of chromosomes 3 (Perepnikhatka *et al*. 1999; Ford *et al*. 2015), 4 (Perepnikhatka *et al*. 1999; Anderson *et al*. 2017), and 5 (Coste *et al*. 2006; Selmecki *et al*. 2006, 2008; Ford *et al*. 2015; Todd *et al*. 2019) have previously been shown to confer increased resistance to fluconazole, hence azole drugs cause both cause a generalized increased rate of aneuploidy as well as selection for specific aneuploidies. Resistance has been attributed to increased gene dosage of drug transporters or their transcriptional activators (*CDR1, CDR2, CZR1*, and *MRR1* on chr3 (Sanglard *et al*. 1995, 1997; Chau *et al*. 2004; Coste *et al*. 2004, 2006; Todd and Selmecki 2020), *TAC1* on chr5 (Selmecki *et al*. 2008)), stress response proteins (*PBS2* on chr3 (Todd and Selmecki 2020); *CGR1* on chr4 (Todd and Selmecki 2020)), and the target enzyme (*ERG11* on chr5 (Chau *et al*. 2004; Selmecki *et al*. 2008)). Furthermore, since many genes are affected by an aneuploidy, non-targeted effects may be more common than with single-gene mutations; for example, chr2 aneuploidy selected under caspofungin exposure confers enhanced survival by different mechanisms to hydroxyurea (Yang *et al*. 2019) and tunicamycin (Yang *et al*. 2021b). In some cases, this may provide an enhanced selective effect for aneuploidy to sweep through a population. In other scenarios, gene overexpression could be selectively disadvantageous (Yang *et al*. 2021a), reducing the potential fitness benefit. Whether consistent posaconazole exposure also selects for beneficial aneuploidy has not been determined.

Here, we conducted the first *in vitro* experimental evolution of eight diverse strains of *C. albicans* in the fungistatic drug posaconazole. We found that increases in drug tolerance to posaconazole were common across strain backgrounds, while increases in drug resistance were rare. We also found that adaptation to posaconazole resulted in widespread cross-tolerance to other azoles and widespread increases in genome size.

## MATERIALS AND METHODS

### Strains and evolution

Eight clinical strains of *C. albicans* that span ancestral fitness to fluconazole (Gerstein and Berman, 2020) were selected (FH1, (Fonzi and Irwin 1993); SC5314, (Lockhart *et al*. 2002); T101 (Odds *et al*. 2007); GC75, P75016, P76055, P78048, and P87, (Wu *et al*. 2007)). Freezer stock was streaked onto YPD agar plates, a standard lab yeast rich medium (2% w/v peptone, 2% w/v yeast extract, 1.8% w/v agar, 1% w/v glucose, 0.00016% w/v adenine sulfate, 0.00008% w/v uridine, 0.1% v/v of each chloramphenicol and ampicillin), and incubated at room temperature for 72 h. Twelve single colonies from each of the eight strains were chosen haphazardly and transferred into 1 mL of liquid YPD in a 3 mL 96-well box, sealed with Breathe-Easier sealing membranes (Electron Microscopy Sciences, PA, United States), and incubated for 24 h at 30 °C, creating twelve replicate lines from each strain. From each replicate, 100 µL was frozen down in 30 % glycerol and stored at −70 °C as the ancestral culture.

We initiated two sets of evolution experiments similarly. To initiate both sets, the optical density was measured from ancestral replicates grown in YPD, and wells were standardized to an OD_600_ of 0.01. A 1:10 dilution was then done in parallel into either YPD + 0.5 μg/mL posaconazole (hereafter referred to as POS) or YPD alone (YPD). For the first set, replicate lines were initially incubated statically at 30 °C for 24 h, followed by 1:1000 serial dilutions into fresh medium (POS or YPD) every 24 h for four days, for a total of ∼50 generations of evolution. The second set of experiments followed a very similar method, except all replicates were initially incubated statically at 30 °C for 72 h, and then four 1:1000 serial dilutions were done into fresh POS medium every 72 h. Twelve replicates from strains P87, GC75, and SC5314 were evolved with both 24 h and 72 h transfers in a POS pilot study that followed the same protocol before the main studies.

In all cases, evolved replicates were frozen down in triplicate in 15 % glycerol after the fifth transfer and stored at −70 °C. In total, 132 replicates were evolved for each transfer duration in POS ((12 replicates x 3 strains) + (12 replicates x 8 strains) = 132 replicates) while 96 replicates were evolved for each in YPD. Replicates were considered extinct at the end of the experiment if they were unable to be revived from the evolved freezer stock.

### Fitness in the evolutionary environment

Fitness in the evolutionary environment was measured as OD_600_ after 24 h and 72 h incubation. Ancestral and evolved replicates were grown from frozen by transferring 5 µL of thawed freezer stock into 500 mL liquid YPD and incubated for 48 h at 30 °C. Each replicate was standardized to OD_600_ 0.01 in liquid YPD. Next, 100 μL of standardized culture was placed into each well of a 96-well round bottom plate, and 100 μL of YPD plus posaconazole was added to each well for a final concentration of 0.5 μg/mL posaconazole. All plates were covered with a Breathe-Easier sealing membrane and incubated statically at 30 °C for 72 h, with OD_600_ measurements taken every 24 h.

### Drug Susceptibility

Drug susceptibility was measured by disk diffusion assays in posaconazole and fluconazole in all replicates. In addition, SC5314 replicates were also assayed in miconazole, clotrimazole, voriconazole, 5-fluorocytosine, and nystatin. Posaconazole disks were prepared by adding 4 μL of 0.625 mg/mL posaconazole stock in DMSO to blank susceptibility disks (Fisher Scientific, Ottawa, ON, Canada) for a final concentration of 2.5 mg. All other susceptibility disks were purchased: fluconazole (25 μg) (Fisher Scientific, Ottawa, ON, Canada); miconazole (50 μg), clotrimazole (50 μg), voriconazole (1 μg), 5-fluorocytosine (1 μg), and nystatin (100 IU) (BioRad Laboratories, Hercules, California, USA).

Ancestral and evolved replicates were grown from frozen in liquid YPD at 30 °C for 48 h. Each replicate was standardized to OD_600_ 0.01. Next, 100 µL of standardized culture was spread, in duplicate, onto 15 mL YPD agar plates, using sterile 5 mm glass beads. A single drug susceptibility disk was applied to the center of each plate, and plates were incubated at 30 °C. After 48 h, each plate was placed on a lightbox and photographed from above using a Canon EOS Rebel SL2.

Photographs were cropped, converted to 8-bit, inverted, and brightness and contrast were altered using ImageJ (Schneider et al., 2012) to obtain bright colonies against a black background. Resistance (RAD_20_) and tolerance (FoG_20_) were quantified from the images using the *diskImageR* R package, following recommendations specified in the diskImageR vignette V2 (Gerstein *et al*. 2016), https://www.microstatslab.ca/diskimager.html). The reported RAD_20_ and FoG_20_ values are averages across multiple biological and technical replicates.

### Ploidy Variation

Flow cytometry was used to determine if evolved replicates had altered ploidy (all were initially diploid). Ancestral and surviving evolved replicates were fixed, stained, and measured in parallel. 5 µL of each replicate was inoculated from frozen into 500 µL of liquid YPD in a deep 96-well box, covered with a Breathe-Easier sealing membrane, and shaken at 350 rpm at 30 °C for 48 h. After 48 h, 10 µL was subcultured into 500 µL of fresh media, and shaken at 350 rpm at 30 °C for 4 h. 200 µL of subculture was then transferred to a 96-well round bottom plate and pelleted. Pellets were resuspended in 20 µL of 50:50 Tris-EDTA (TE), fixed by slowly adding 180 µL of 100 % cold ethanol, and stored wrapped in aluminum foil at −20 °C for at least 12 h.

The fixed culture was pelleted, washed in 200 µL of TE, pelleted again, and resuspended in 50 µL of 1 mg/mL RNAse A solution (New England Biolabs, Ipswich, Massachusetts, United States) and statically incubated at 37 °C for 3 h. After the 3 h incubation, the replicates were pelleted and resuspended in 50 µL TE and 50 µL of 1:100 SYTOX: TE solution and incubated at room temperature in the dark overnight. The next day the replicates were pelleted then resuspended in 700 µL of TE. All centrifugation steps were done at 1000 x g for 5 minutes.

Flow cytometry was performed on an SH800S Cell Sorter (Sony Biotechnology Inc, San Jose, California, United States). All replicates had an event rate of 600-1000 events/second and a total of 10 000 events were recorded. Data was uploaded to FlowJo (Tree Star, Ashland, Oregon, United States), and debris was excluded via gating. Each replicate population was then fit with the Watson (pragmatic) cell cycle algorithm (Watson *et al*. 1987) to determine the mean G1 peak.

### DNA extraction

Genomic DNA was extracted from two ancestral strain replicates, and 12 evolved replicates from SC5314, the strain used for the *C. albicans* reference genome. 30 μL of each replicate were transferred into 3 mL of YPD and incubated shaking overnight at 37 °C. The culture was centrifuged at 2 500 rpm for 3 minutes, and the supernatant was discarded. The pellet was resuspended in 500 µL of TENTS buffer (100 mM NaCl, 10 mM Tris pH 8.0, 1 mM EDTA, 2% Triton X 100, and 1% SDS), 100 µL of glass beads, and 200 µL phenol:chloroform: IAA and vortexed for 20 minutes at 4 °C followed by centrifugation for 10 minutes at 13 500 rpm. After centrifugation, 350 µL of supernatant was transferred to a sterile microcentrifuge tube, and 1 mL of cold 100 % ethanol was added and left overnight at −20 °C to allow for DNA precipitation.

This was followed by centrifugation at 13 500 rpm for 10 minutes, and the pellet was resuspended in 50 µL of molecular biology grade water. One microliter of 10 mg/mL RNAse A was added and incubated at 37 °C for 1 h, after which 2 µL of 20 mg/mL proteinase K (Fisher Scientific, Ottawa, Ontario, Canada) was added and incubated at 37 °C for 1.5 h. After incubation, 200 µL of molecular biology grade water and 300 µL of phenol:chloroform: IAA was added, and samples were centrifuged at 13 500 rpm for 10 minutes. Next, the supernatant was transferred to a sterile microcentrifuge tube, 4 µL of 5 M NaCl and 400 µL of cold 100 % ethanol were added and DNA was precipitated for at least 10 minutes at −20 °C. Tubes were then centrifuged at 13 500 rpm for 10 minutes, the supernatant was discarded, and the pellet was left overnight. The DNA pellet was then resuspended in 40 µL of molecular biology grade water.

DNA quality was assessed using the Thermo Scientific™ NanoDrop 2000, and DNA concentration was measured using a Qubit® 2.0 Fluorometer (following the Invitrogen™ Qubit™ dsDNA BR Assay Kit) (Thermo Fisher Scientific, Waltham, Massachusetts, United States). Whole-genome sequencing was performed by the Microbial Genome Sequencing Center (Pittsburgh, USA) using the Illumina NextSeq 550 platform to a calculated depth of ∼40× pairedend reads of 2 × 150 bp.

### Karyotype analysis

To analyze the whole-genome sequences for karyotypic variation in evolved SC5314 replicates, we used Y_MAP_, a computational pipeline that visualizes copy number variation and loss of heterozygosity (Abbey *et al*. 2014). Mitochondrial DNA was excluded from the coverage analysis. Paired-end read data were uploaded and analyzed using the SC5314 A21-s02-m09-r07 reference genome. Both baseline and experimental ploidy were left at the default values (two, i.e., diploid), and correction was enabled for GC-content bias and chromosome-end bias.

Smaller-scale CNVs were also evaluated by comparing the copy number at each position in evolved strains to the copy number in two sequenced ancestral strains. Positions elevated only in evolved strains were further examined individually, and the location mapped to the genome using the Candida Genome Database (candidagenome.org).

## RESULTS

### Survival and fitness in posaconazole evolution

Replicate lines from eight clinical strains of *C. albicans* were passaged with 1:1000 dilutions every 24 h or 72 h, for a total of five transfers in YPD + 0.5 ug/mL posaconazole (POS) and in YPD. The level of posaconazole was above the MIC_50_ of all ancestral strains, representing a strong selective pressure. We use the term “ancestral replicates”to indicate populations initiated from single colony replicates before evolution, and the term “evolved replicates”to refer to the replicates that survived the evolution experiment. No replicates survived in the POS 24 h transfer experiment, while approximately half of the replicates (67) survived to the end of the POS 72 h transfer experiment. Replicate survival to 72 h transfers in POS was not equal among strain backgrounds (Figure 1). Surprisingly, ancestral fitness in the evolutionary drug environment was not significantly correlated with the number of surviving replicates after evolution (Figure 1; ancestral fitness measured as optical density after 72 h of growth in POS, Pearson’s correlation, t_6_ = 0.91, *P* = 0.40). This lack of correlation indicates that strain-specific genomic differences that influence something other than ancestral fitness likely correlate with evolvability to POS. All replicate lines survived 24 h and 72 h transfers in the standard rich medium YPD. The majority of the remaining analysis is based on the 72 h transfer experiment in POS, except where indicated.

**Figure 1.**
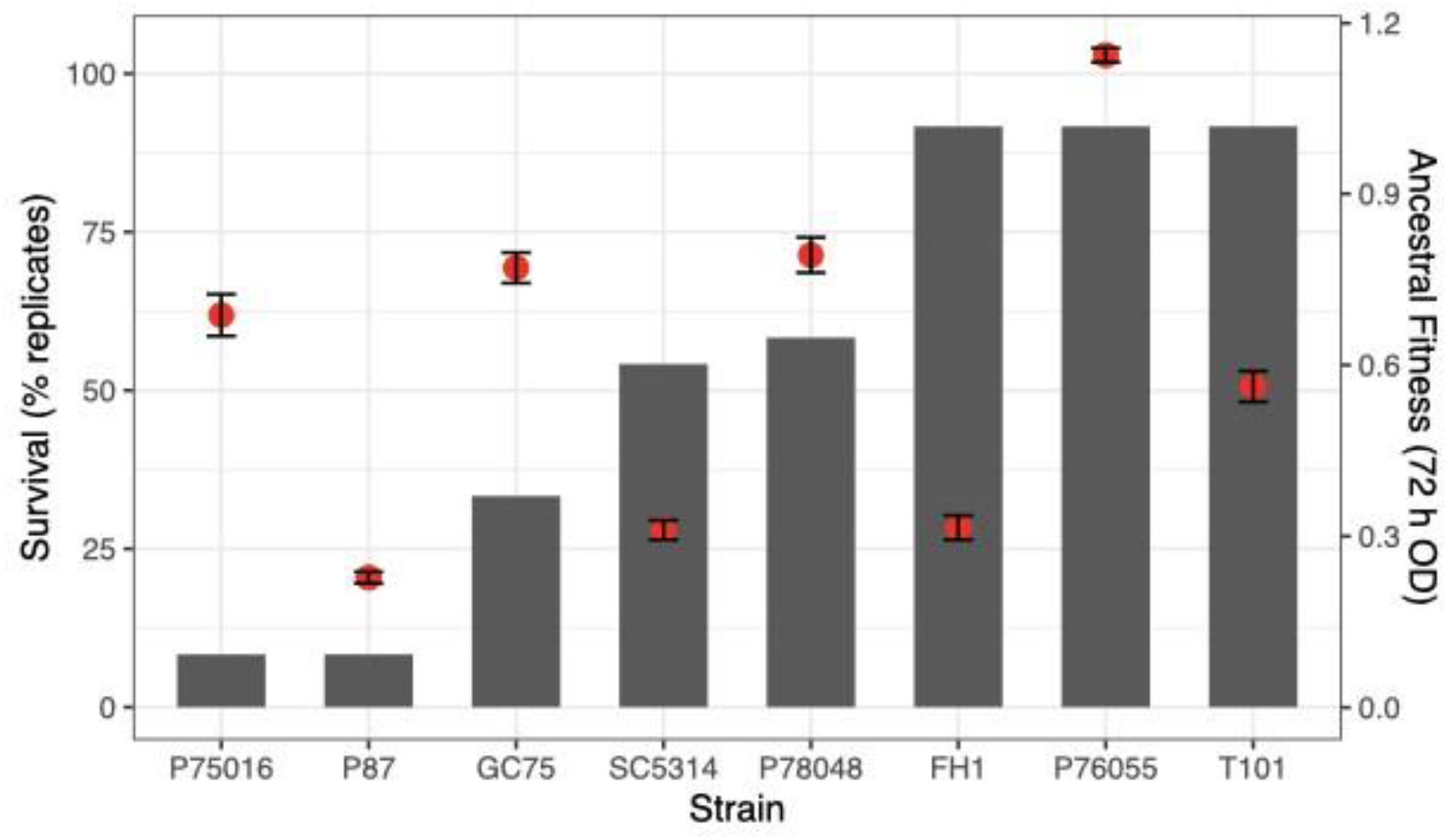
Differential survival of evolved replicates to five passages posaconazole. The grey bars indicate the percentage of replicates that survived to the end. Red dots indicate ancestral fitness (optical density after 72 h of growth in the evolutionary environment), shown on the right vertical axis. Each dot is the mean +/- SE of two biological replicates. The percent of replicates that survived was associated with ancestral strain fitness.

Evolved replicates from the five strains with the highest number of surviving replicates (P78048, SC5314, FH1, P76055, T101) grew to a higher optical density than ancestral replicates after 24 h in POS (Figure 2; t-test results in Table 1) and four of the five strains retained the advantage after 72 h of growth (Figure 2, Table 1). There were not enough surviving replicates from P87 or P75016 to properly conduct statistical tests, though the one surviving replicate from P75016 also very clearly has a fitness advantage over the ancestral replicates. Evolved replicates from GC75 had no improvement over ancestral replicates at 24 h or 72 h. Across all strain backgrounds, there was an inconsistent tradeoff between improvement in fitness in drug and reduction of fitness in YPD (Figure S1; t-test results in Table 2). Evolved replicates from three strains (P78048, SC5314, FH1) had a minor reduction in fitness in YPD at 24 h, which continued to 72 h for SC5314 and FH1. T101 had a minor improvement in fitness at 24 h in YPD. Of note, the magnitude of fitness difference between statistically-significant ancestral and evolved replicates tended to be considerably less in YPD compared to drug (Table 1).

**Table 1.** T-test results comparing optical density of ancestral and evolved replicates grown for 24 h (left) and 72 h (right) in POS (top) and YPD (bottom).

| Strain | Posaconazole 24 h |  | Posaconazole 72 h |  |
| --- | --- | --- | --- | --- |
|  | Evol-Anc | statistic | Evol-Anc | statistic |
| GC75 | 0.08 | $t_{7.9} = -2.0, p = 0.09$ | -0.08 | $t_{10.8} = 0.8, p = 0.45$ |
| SC5314 | 0.17 | $t_{12.2} = -2.9, p = 0.013$ | 0.18 | $t_{13.6} = -2.2, p = 0.048$ |
| P78048 | 0.5 | $t_{8.2} = -5.4, p < 0.001$ | 0.33 | $t_{15.0} = -3.9, p = 0.002$ |
| FH1 | 0.13 | $t_{11.5} = -11.4, p < 0.001$ | 0.08 | $t_{19.8} = -4.3, p < 0.001$ |
| P76055 | 0.2 | $t_{13.5} = -2.9, p = 0.013$ | 0.09 | $t_{15.0} = -3.0, p = 0.009$ |
| T101 | 0.12 | $t_{13.2} = -2.3, p = 0.036$ | 0.09 | $t_{20.1} = -1.5, p = 0.15$ |

| Strain | YPD 24 h |  | YPD 72 h |  |
| --- | --- | --- | --- | --- |
|  | Evol-Anc | statistic | Evol-Anc | statistic |
| GC75 | -0.15 | $t_7 = -1.65, p = 0.14$ | -0.12 | $t_7 = 1.36, p = 0.22$ |
| SC5314 | -0.24 | $t_{12.3} = 2.86, p = 0.014$ | -0.24 | $t_{12.3} = 2.74, p = 0.018$ |
| P78048 | -0.09 | $t_{7.2} = -3.5.6, p = 0.009$ | -0.01 | $t_{13.9} = 0.84, p = 0.42$ |
| FH1 | -0.08 | $t_{10.6} = 4.25, p = 0.002$ | -0.09 | $t_{11} = 9.05, p < 0.001$ |
| P76055 | 0.02 | $t_{11.0} = -1.3.6, p = 0.22$ | -0.002 | $t_{11.2} = 0.14, p = 0.89$ |
| T101 | 0.04 | $t_{20.6} = -3.0.6, p = 0.007$ | 0.01 | $t_{20.7} = -1.16, p = 0.26$ |

**Figure 2.**
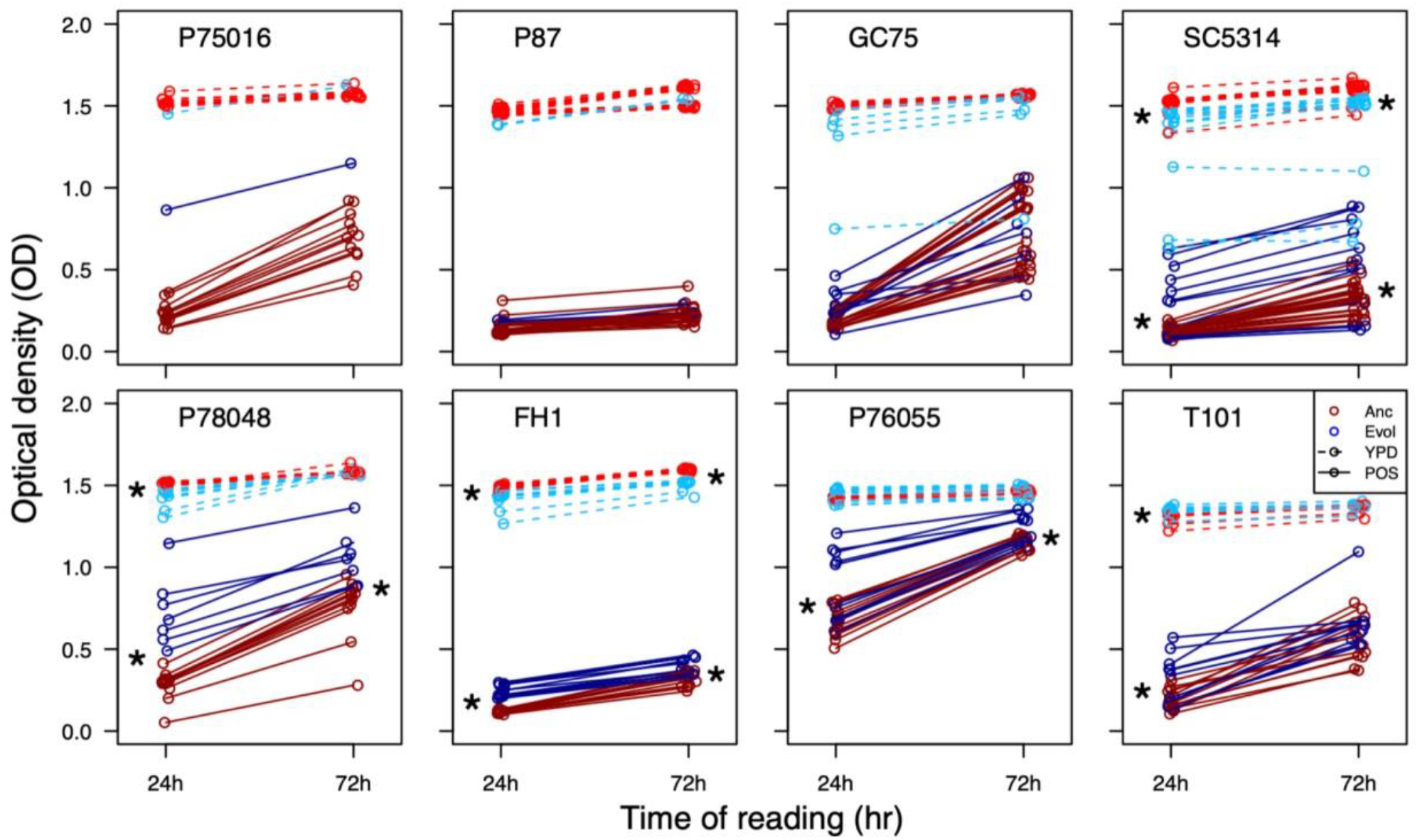
Fitness of ancestral and evolved replicates grown in YPD and POS. Optical density was measured at 24 h and 72 h. Shown here is only the replicates that were evolved through 72 h transfers in POS. Evolved replicates grown in YPD are indicated with brighter colours and dashed lines, replicates grown in POS are darker colours and full lines. Stars indicate statistical significance in a t-test comparing ancestral and evolved replicates at that time point (24 or 72 h of growth).

### Widespread decreases in drug resistance and increases in drug tolerance in POS evolved replicates

Drug susceptibility was computationally quantified as the radius of inhibition on a disk diffusion plate (Gerstein *et al*. 2016). Six strain backgrounds evolved an increase in susceptibility (Figure 3; GC75: t_7.6_ = −2.7, p = 0.029; SC5314: t_12.7_ = −2.25, p = 0.043; P78048: t_6.8_ = −7.7, p = 0.0001; FH1: t_13.0_ = −4.3, p = 0.0008; strains P75016 and P87 had too few replicates for statistical testing). Six replicates from four strains did deviate from the rest and acquired increased resistance (reduced susceptibility; P75016, GC75, SC5314, and P78048). Evolved replicates from the two strain backgrounds with the highest number of surviving replicates did not change in susceptibility (P76065: t_13.6_ = −1.4, p = 0.17; T101: t_15.4_ = 0.8, p = 0.42). The opposite result was seen for drug tolerance, which is computationally determined as FoG_20_, the fraction of growth between the disk and RAD_20_ (Gerstein *et al*. 2016); Figure 3). Evolved replicates from six strain backgrounds increased in tolerance, while two of the strains with a higher number of surviving replicates did not change (Figure 3; GC75: t_11.6_ = −14.5, p < 0.0001; SC5314: t_14.0_ = 12.63, p < 0.0001; P78048: t_6.1_ = −7.3, p = 0.0003; FH1: t_19.8_ = 0.6, p = 0.59; P76055: t_18.0_ = −2.2, p = 0.042; T101: t_20.1_ = 0.9, p = 0.50). Evolved replicates from across strain backgrounds more consistently increased in tolerance rather than resistance, suggesting that tolerance is either more evolvable than resistance, or was the primary phenotype under selection in POS.

**Figure 3.**
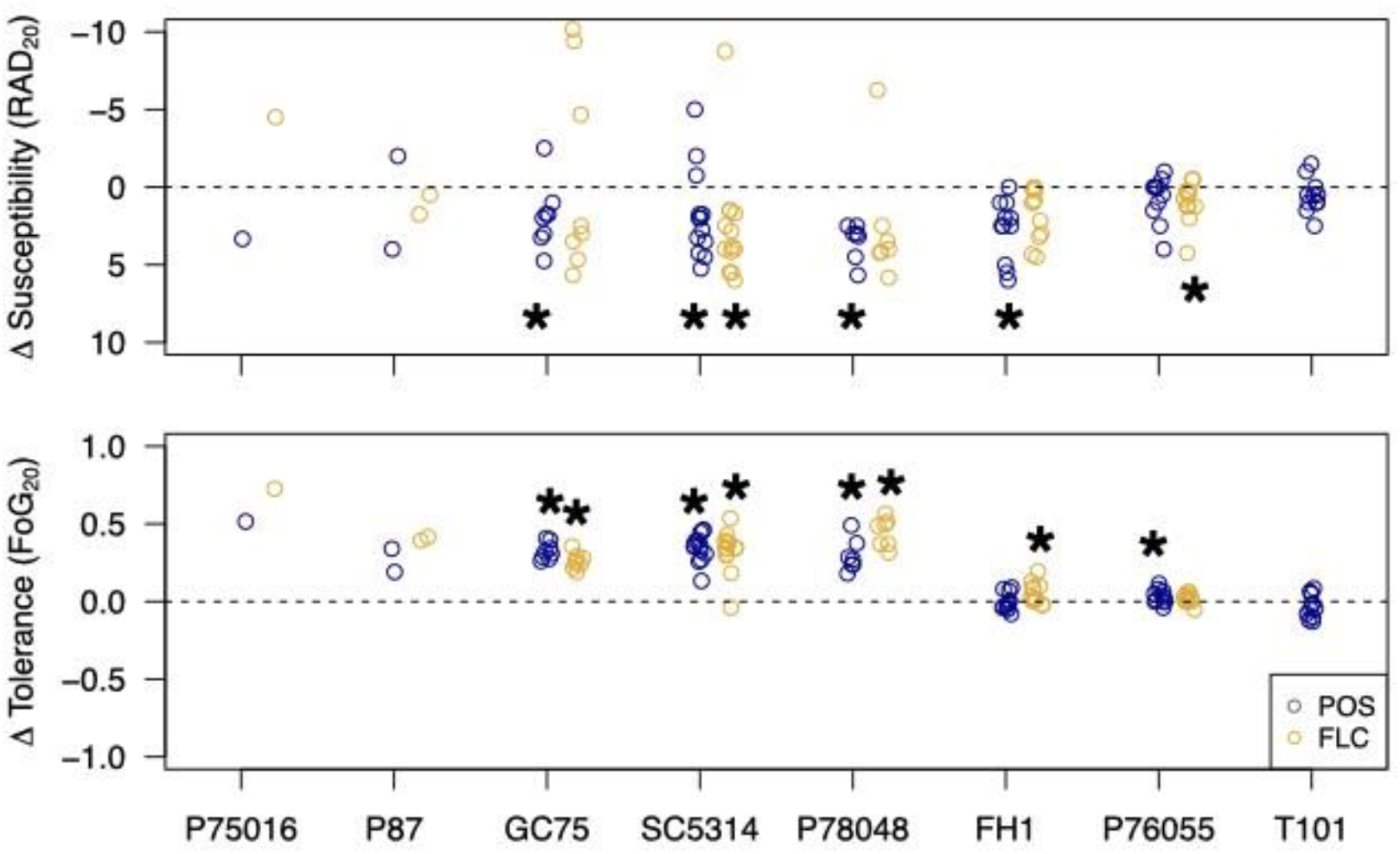
Susceptibility (top) and tolerance (bottom) of posaconazole-evolved replicates assayed on posaconzolc and fluconazole disks. Shown is the difference in phenotype between the evolved replicate and the median of 12 ancestral replicates. A negative change in susceptibility in the evolved replicate indicates an increase in resistance, and the y-axis of the top panel is reversed to reflect this. Stars indicate a significance difference compared to the ancestral replicates from a t-test (p < 0.05).

All evolved replicates were also assayed for the evolution of cross-resistance and/or cross-tolerance to the most common triazole, fluconazole (FLC). Results in FLC were similar to POS; evolved replicates tended to increase in FLC susceptibility (decrease in resistance) yet increase in FLC tolerance. Interestingly, in a number of strain backgrounds, POS-evolved replicates increased in FLC resistance to a higher degree than they had in POS. This lead on occasion to the loss of a significant difference in susceptibility between ancestral and evolved replicates in some strain backgrounds, though it is clear that the majority of evolved replicates in most backgrounds increased in FLC susceptibility (Figure 3; FLC resistance—GC75: t_7.1_ = −0.52, p = 0.62; SC5314: t_13.7_ = −2.51, p = 0.025; P78048: t_6.0_ = −0.93, p = 0.39; FH1: t_11.2_ = −1.9, p = 0.08; P76055: t_16.8_ = −2.6, p = 0.02; tolerance—GC75: t_13.2_ = −13.2, p < 0.0001; SC5314: t_13.5_ = −8.26, p < 0.0001; P78048: t_6.0_ = −12.7, p < 0.0001; FH1: t_11.2_ = −1.9, p = 0.09; P76055: t_18.8_ = 1.3, p = 0.21). The ancestral and evolved replicates of T101 are resistant to FLC and changes to FLC tolerance in evolved replicates can not be examined in this framework.

### Inconsistent changes in drug resistance and drug tolerance in YPD evolved replicates

Replicates that were evolved with 24 h and 72 h transfers in YPD exhibited much smaller changes in resistance and tolerance, though some evolved strains did differ significantly compared to the ancestors (Figure S1, Table S1). There was no apparent pattern between strain background and the specific evolved changes. Evolved replicates from strain P87 (a strain with few surviving POS replicates) in both the 24 h and 72 h transfer experiments had significantly increased susceptibility. By contrast, evolved replicates from strain P76055 and T101, the two strains with the most surviving POS replicates, had significantly decreased susceptibility (in both experiments for strain P76055 while only in the 24 h transfers for strain T101). Evolved replicates from strain GC75 in the 24 h transfer experiment decreased in tolerance, while replicates from strain FH1 in the 24 h transfer experiment increased in tolerance.

### Widespread karyotypic changes after posaconazole evolution

Genome size changes in POS evolved replicates were examined with flow cytometry. Evolved replicates from the three strains with the highest proportion of surviving replicates (FH1, P76055, and T101) tended to retain genome sizes similar to the diploid ancestors, while the majority of evolved replicates from other strains varied in genome size (Figure 4). Measured changes in genome size (G1 means) were generally consistent with aneuploidy rather than whole shifts in ploidy (Figure 4, inset panels).

**Figure 4.**
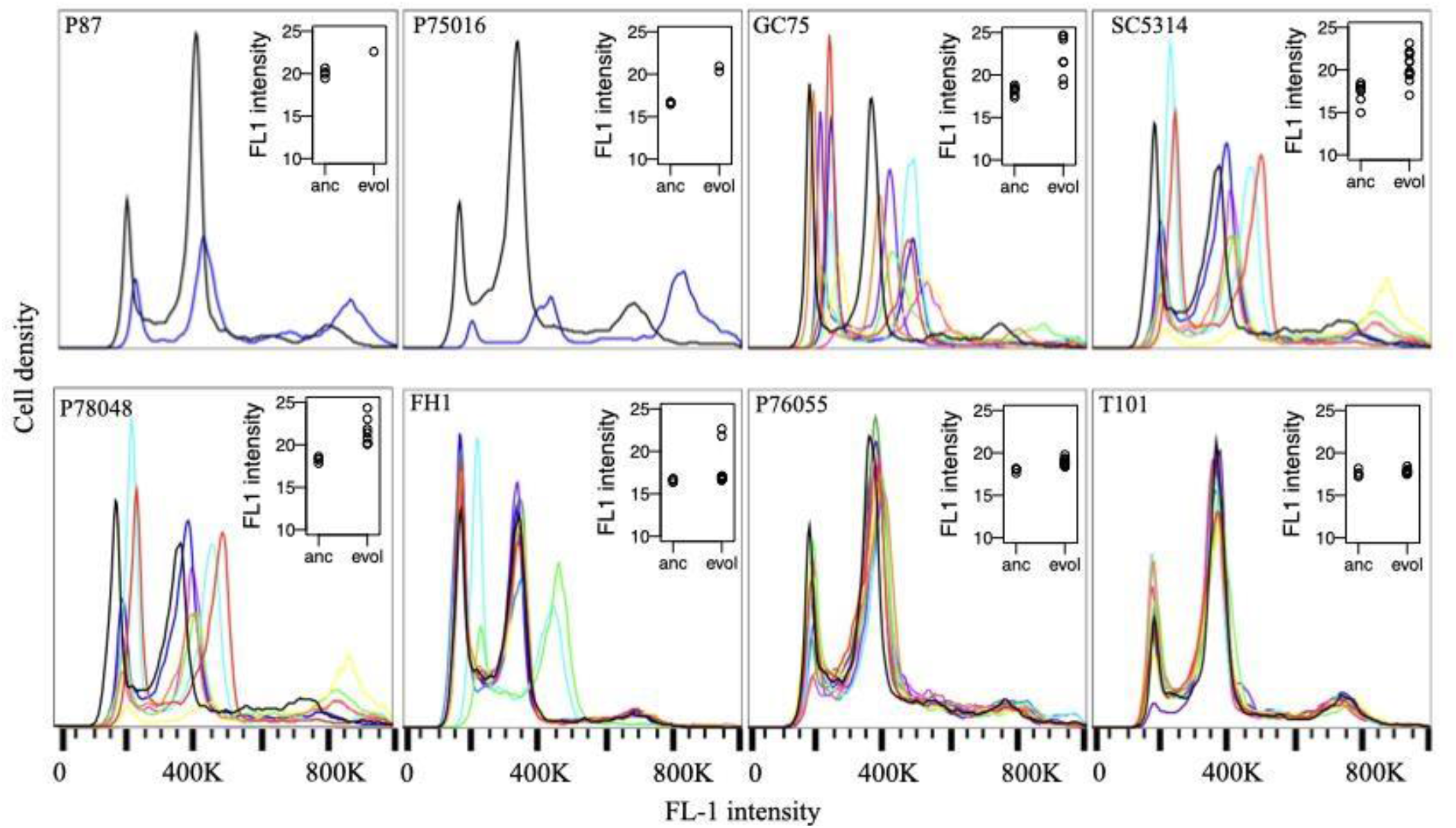
Flow cytometry traces from POS evolved replicates. Each trace is a single evolved replicate, the black trace in each panel is a diploid ancestral replicate. Inset panels display the mean G1 peak from each ancestral (anc) and evolved (evol) replicate for each strain.

### Chr6 and ChrR aneuploidy and cross-azole tolerance in evolved replicates

To examine actual karyotypic changes and how they correlate with drug resistance and tolerance evolution, we performed whole-genome sequencing on 12 evolved replicates from SC5314. All but one evolved replicate had at least one trisomic chromosome. Trisomy of chrR was the most common, with nine replicates having either whole or partial trisomy of this chromosome. Of these nine, two replicates had additional aneuploidies in either chr4 or chr4 and chr6. Two other evolved replicates also had an extra copy of chr6, one alone and one in tandem with an extra copy of chr3 (Figure 5A). We saw no apparent bias towards acquiring an extra copy of the A haplotype or the B haplotype. Most trisomies seem to have swept through the evolved populations, evidenced by a copy number of three from population-level sequencing (Figure 5B). In several cases, the measured copy number was between two and three, likely indicating a polymorphic population, where some cells remained diploid (though we can not rule out that some aneuploidies may be unstable and lost during the grow up from frozen evolved culture for sequencing). A small number of localized copy number variation was also present in evolved lines, and none were suggestive of an adaptive benefit (Table S2). All small CNVs were either associated with long terminal repeats, major repeat sequences, telomeres, or existed as multiple sites on a single aneuploid chromosome in a single background, suggestive of noisy sequencing.

**Figure 5.**
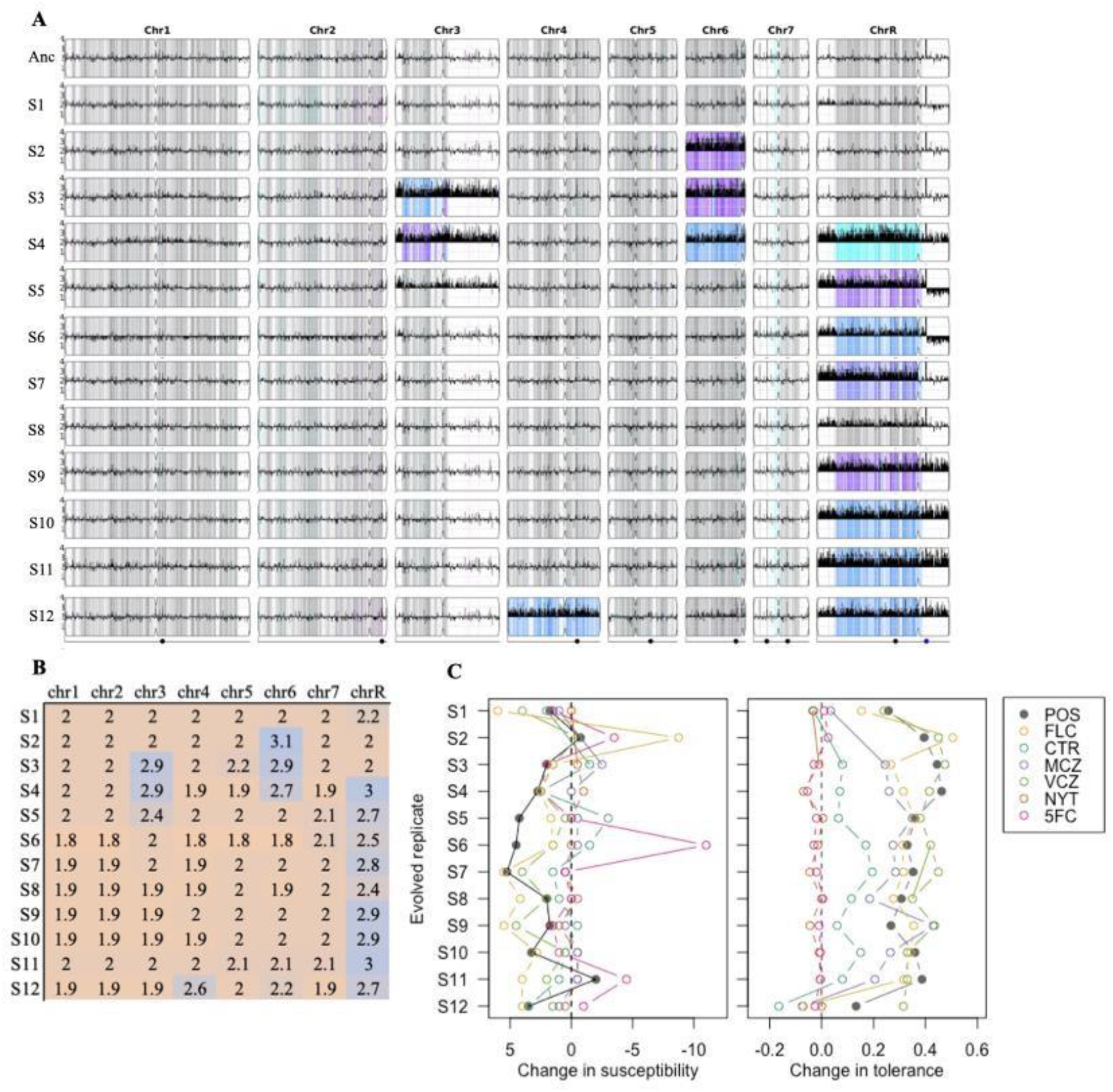
Copy number variation, cross-resistance and cross-tolerance in whole-genome sequenced replicates of SC5314. A) Chromosomal aneuploidy and loss of heterozygosity (LOH) in evolved strains from the Yeast Mapping Analysis Pipeline (YMAP). All strains were compared to the SC5314 reference background. The density of heterozygous SNPs is presented as vertical lines spanning the height of each chromosome cartoon, with the intensity of representing the number of single nucleotide polymorphisms (SNPs) in each 5kb bin. SNPs that remain heterozygous are displayed in grey, while those that have become homozygous (i.e LOH regions) are displayed in the color assigned to the homolog that is retained (cyan for the ‘A’ allele, magenta for the ‘B’ allele, blue for AAB and purple for ABB). The white regions represent ancestral LOH blocks. CNVs per position are displayed as black histograms drawn vertically from the centerline along the length of the chromosome cartoon.The numbers on the Y axis represent the relative chromosome copy numbers, based upon the whole genome ploidy. The centromere loci are illustrated as an indentation in the scaled chromosome cartoon. The dots represent the positions of the major repeat sequences. B) Median copy number of all reads mapped to each chromosoms. C) The change in susceptibility (left) and tolerance (right) in evolved replicates relative to the SC5314 ancestor. Susceptibility and tolerance were measured from disk assays for posaconazole (POS), fluconazole (FLC), clotrimazole (CLT), miconazole (MCZ), voriconazole (VCZ), nystatin (NYT) and 5-fluorocytosine (5FC). A negative change in susceptibility is an increase in resistance.

We also subjected these 12 evolved SC5314 replicates to a panel of five additional antifungal drugs to look for cross-resistance and cross-tolerance: voriconazole (VCZ), a triazole like POS and FLC; clotrimazole (CTR) and miconazole (MCZ), which are imidazoles; nystatin (NYT), a polyene; and 5-fluorocytosine (5-FC). Similar to what we found across strain backgrounds for POS and FLC, very few replicates exhibited decreased susceptibility to any additional drugs (Figure 5B). By contrast, nearly all evolved replicates showed increased tolerance to all five azole drugs examined and essentially no change in NYT or 5-FC. There was no clear phenotypic pattern to differentiate evolved strains with ChrR trisomy (or other trisomies) from other replicates, indicating that multiple genotypic pathways seem to provide increased tolerance to POS and that cross-tolerance to other azoles is likely to be a common feature of adaptation to POS.

## DISCUSSION

Experimental laboratory evolution has been an effective method to study pathways of adaptation in a variety of biological contexts (Kawecki *et al*. 2012; Cooper 2018). Laboratory evolution of microbes at high population sizes (at times termed ‘experimental evolution’ or ‘adaptive laboratory evolution’) is a particularly powerful way to examine parallelism and constraint in evolution at genomic and phenotypic levels (Cooper 2018; Gerstein and Sharp 2021). We evolved 12 replicates from eight different strain backgrounds of the opportunistic human fungal pathogen *Candida albicans* to 0.5 ug/mL of the drug posaconazole (POS), with transfers every 24 h or 72 h. This level of drug exceeds where we find robust growth in the absence of stress from most ancestral strains, and is an order of magnitude above the defined epidemiological cutoff value of resistance (Arendrup *et al*. 2011; Procop 2020). As drug tolerance is defined as slow growth in the presence of high levels of drug, we reasoned that a more extended transfer period might select for drug tolerance to evolve. Unfortunately, none of the evolved replicates survived the 24 h transfers. In the 72 h transfer experiment, we also found high extinction (∼50% of the replicates), yet the majority of surviving replicates indeed increased in POS tolerance and acquired cross-tolerance to FLC and other azole drugs. Very few studies have (yet) directly examined how different conditions influence how readily tolerance evolves in *C. albicans*. In an experiment where replicates were evolved with 72 h transfers to a sub-inhibitory level of fluconazole (i.e., a level of FLC below the MIC), changes in tolerance were also common, but both increases and decreases were observed (Gerstein and Berman 2020). We found a similar result here in YPD when replicates were transferred for 24 h: replicates from one strain background significantly increased in tolerance while replicates from a second significantly decreased. Additional experiments are required to tease apart how different drugs, the level of drug stress (sub-inhibitory vs. inhibitory), and other specifics of environmental exposure drive differences in the propensity to acquire increases in drug tolerance. Nevertheless, it seems clear that there is potential for drug tolerance to evolve quickly, which may have important clinical implications, particularly in cases of persistent candidemia infections when populations are exposed to drug stress for long periods of time, yet the underlying strain remains drugsusceptible (Rosenberg *et al*. 2018).

Evolved POS replicates from the majority, but not all, strain backgrounds increased in their growth ability in the evolutionary level of drug. However, this improvement was primarily restricted to the specific drug level strains were evolved in, and the improvement in most replicates was relatively low. Consistent with these results in POS, very few replicates evolved to a subinhibitory level of FLC increased in FLC resistance (Gerstein and Berman 2020), though a second evolution experiment done at the same sub-inhibitory level of FLC found ∼30% of replicates had increased resistance (Todd and Selmecki 2020). An evolution experiment that similarly paired a strong selective pressure with a long period of time (7 days) between transfers to caspofungin, a different drug class, also found that many evolved diploid lines did not increase in resistance by the end of the 59-day experiment (Avramovska *et al*. 2021). In contrast, increased FLC MIC was evident in the earliest *C. albicans* evolution experiments conducted in the T101 background in continually increasing levels of FLC for ∼330 generations (Cowen *et al*. 2000).

Changes in genome size were also widely observed in our POS evolved replicates, similar to previous *in vitro* evolution experiments in FLC (Selmecki *et al*. 2006, 2009; Coste *et al*. 2007; Ford *et al*. 2015; Gerstein and Berman 2020; Selmecki *et al*. unpublished) and in strains passaged through a mouse GI model (Ene *et al*. 2018). Aneuploidies identified in FLCevolved strains typically involve genes known to be involved in FLC resistance mechanisms such as Hsp90, efflux pumps, and multidrug transporters *MRR1, CDR1, CDR2, CRZ1* on chr3 (Mount *et al*. 2018; Todd and Selmecki 2020), and *ERG11, TAC1*, and calcineurin genes on chr5 (Selmecki *et al*. 2008). In contrast, in our SC5314 evolved isolates, we predominantly found extra copies of chrR (9 isolates), chr6 (3 isolates), and chr3 (two isolates, in tandem with chr6 aneuploidy in both cases). These aneuploidies were concurrent with increases in tolerance to POS (and other azoles), but not increases in resistance to POS or other tested drugs classes. This is somewhat surprising, as a previous study in a different strain background found that an extra copy of chrR increased resistance to FLC, ketoconazole, and miconazole (Li *et al*. 2015). Chromosome 3 aneuploidy was previously identified to confer an increase in FLC tolerance, due at least in part to an extra copy of the urea transporter *NPR2* (Mount *et al*. 2018). Interestingly, chr6 and chrR were previously found to be the two most common aneuploidies remaining in otherwise euploid strains after 28 days of daily passaging of initially-tetraploids and initiallyaneuploid strains in standard lab YPD (Hickman *et al*. 2015). This suggests that either these chromosomes carry genes that are beneficial in the context of *in vitro* evolution in general, or perhaps they simply carry the lowest cost in an otherwise euploid background in an environment where aneuploidy frequently occurs.

The rapid evolution of tolerance and cross-tolerance through *in vitro* evolution has not been previously observed in azoles nor linked to aneuploidy. Todd and Selmecki (2020) did, however, identify three isolates that evolved in an FLC *in vitro* experiment that had increased copy number of a subset of efflux pump, multidrug transporters, and stress response genes on chr3, leading to increased tolerance and resistance to several azoles. Although the mechanism(s) that underlie drug tolerance are still being resolved, there are putative tolerance-associated genes on chrR to target for follow-up studies. Transcriptomics and phenotypic analyses demonstrated that all Rim proteins (including *RIM9*, on chrR) were important for FLC tolerance (Garnaud et al. 2018). Also intriguing and worthy of further study is *GZF3*, a GATA-type transcription factor of unknown function, one of two genes (alongside *CRZ1*) identified in an overexpression screen of 572 genes for fluconazole tolerance (Delarze *et al*. 2020). Tolerance has both a genetic component, likely involving genes in membrane biosynthesis/integrity and the stress response pathways (Cowen *et al*. 2014; Rosenberg *et al*. 2018; Berman and Krysan 2020; Todd and Selmecki 2020), and an environmental component, as growth conditions have been linked to the degree of tolerance exhibited (Gerstein *et al*. 2016; Berman and Krysan 2020). POS is not a substrate for *MDR1* or *FLU1* encoded efflux pumps (Chau *et al*. 2004; Hof 2006), hence there are likely to be both pan-azole and azole-specific genetic mechanisms underlying this complex trait.

Experimental evolution studies have contributed a wealth of knowledge towards understanding a wide range of factors that influence evolutionary dynamics. We used an *in vitro* evolution framework at a high level of the second-generation azole drug posaconazole to evolve replicates from diverse *C. albicans* strains. We found that the survival of replicates was strongly dependent on strain background. The majority of evolved replicates improved in rapid growth in the evolutionary level of drug, yet very few replicates increased in POS resistance. Replicates from strains with fewer surviving replicates were more likely to increase in POS drug tolerance and more likely to increase in genome size. Three chromosomal aneuploidies were observed in parallel in multiple evolved lines; these are largely different from those that have been shown to confer an increase in resistance to the related azole drug fluconazole, indicating that the genetic pathway to acquiring posaconazole tolerance is distinct. As the azole drugs are fungistatic rather than fungicidal, further work is required to determine whether increases in tolerance upon repeated exposure to drugs are common in a clinical setting and whether tolerance is a stepping-stone in the path to resistance or represents a distinct peak in an adaptive trajectory.

## DATA AVAILABILITY

Raw data and R scripts used for statistical analyses and to generate figures is available at https://github.com/acgerstein/posaconazole-evolution

## ACKNOWLEDGEMENTS

We thank Ola Salama for feedback on troubleshooting experiments and helping shake endless disk assays. We appreciate Dr. Ayush Kumar, Dr. Peter Pelka and Dr. Silvia Cardona for use of plate readers.

## FUNDING

ACG was funded by an NSERC Discovery Grant, Start-up funding from the University of Manitoba and a University of Manitoba University Research Grants Program grant. RJK and BS were supported by a University of Manitoba Faculty of Science Undergraduate Research Award, QW and RS were supported by NSERC Undergraduate Student Research Awards.

## SUPPORTING MATERIAL

**Table S1.** Two-way ANOVAs comparing drug susceptibility and drug tolerance among ancestral replicates, replicates evolved in YPD with 24 h transfers and replicates evolved in YPD with 72 h transfers.

| Strain | Susceptibility ANOVA | Tolerance ANOVA |
| --- | --- | --- |
| P75016 | $F_{2, 32} = 0.02, p = 0.98$ | $F_{2, 32} = 0.37, p = 0.70$ |
| P87 | $F_{2, 32} = 21.14, p < 0.0001$ | $F_{2, 32} = 1.61, p = 0.22$ |
| GC75 | $F_{2, 32} = 1.87, p = 0.17$ | $F_{2, 32} = 10.73, p = 0.0003$ |
| SC5314 | $F_{2, 32} = 0.05, p = 0.95$ | $F_{2, 32} = 0.41, p = 0.67$ |
| P78048 | $F_{2, 32} = 1.00, p = 0.381$ | $F_{2, 32} = 0.46, p = 0.63$ |
| FH1 | $F_{2, 32} = 1.74, p = 0.19$ | $F_{2, 32} = 10.97, p = 0.0003$ |
| P76055 | $F_{2, 32} = 12.47, p < 0.0001$ | $F_{2, 32} = 2.34, p = 0.11$ |
| T101 | $F_{2, 32} = 10.82, p = 0.0002$ | $F_{2, 32} = 3.56, p = 0.04$ |

**Figure S1.**
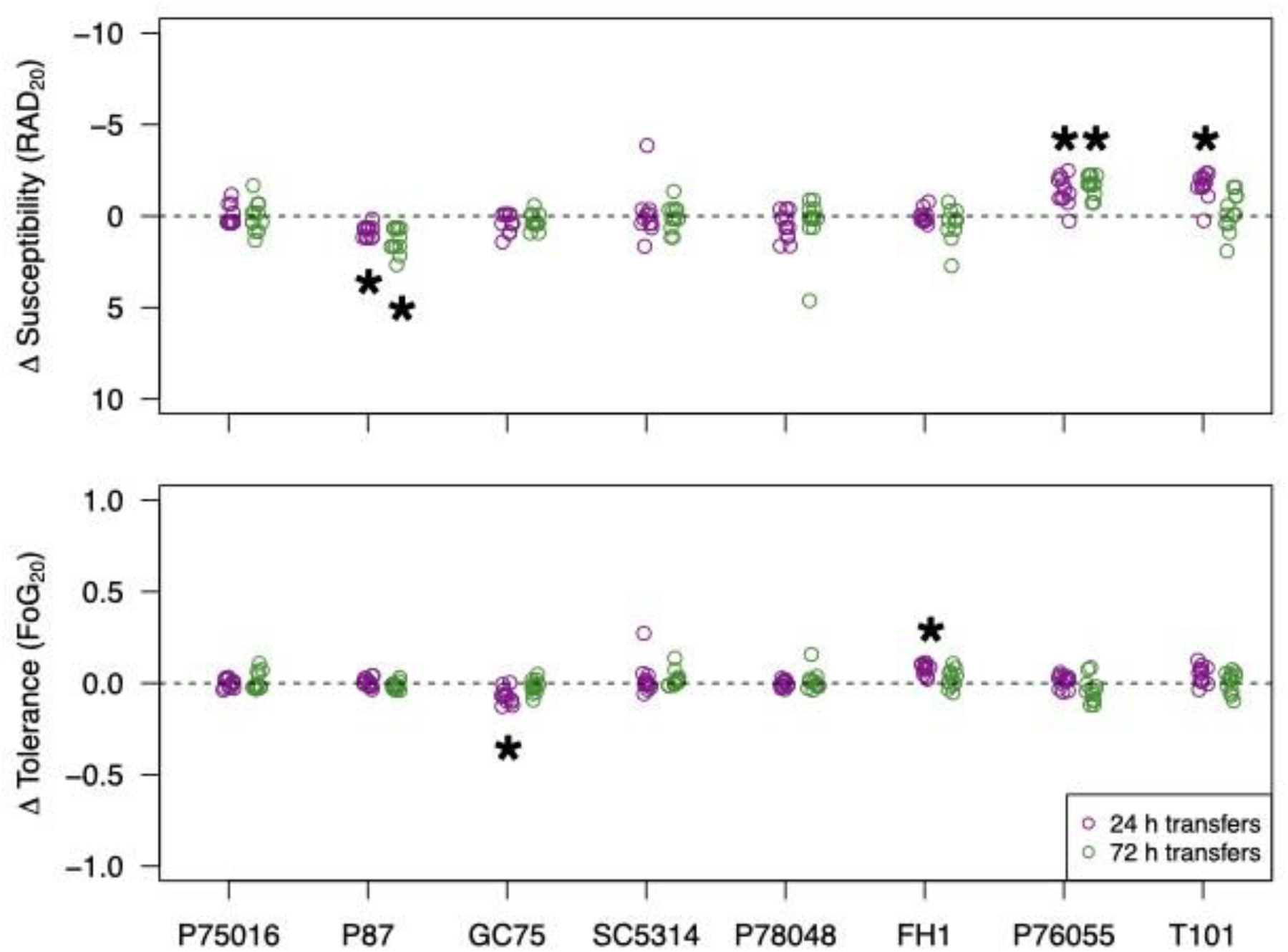
Susceptibility (top) and tolerance (bottom) from YPD-evolved replicates from 24 h (purple dots) and 72 h (green dots) transfer experiments assayed on posaconzolc disks. Shown is the difference in phenotype between the evolved replicate and the median of 12 ancestral replicates from each. A negative change in susceptibility in the evolved replicate indicates an increase in resistance, and y axis of the top panel is reversed to reflect this. Stars indicate a significance difference compared to the ancestral replicates from a post-hoc Tukey test following a significant ANOA (p < 0.05).

## Notes

https://github.com/acgerstein/posaconazole-evolution

### Competing Interest Statement

The authors have declared no competing interest.

